# General aspects of visceral leishmaniasis with or without HIV co-infection in Northeast Brazil

**DOI:** 10.1101/403683

**Authors:** Uiara Regina Silva de Lima, Luciano Vanolli, Elizabeth Coelho Moraes, Jorim Severino Ithamar, e Conceição de Maria Pedrozo e Silva de Azevedo

## Abstract

**Background:** The case of visceral leishmaniasis (VL) and human immunodeficiency virus (HIV) co-infection was reported in Spain first in 1985, and coincidence of these diseases has also been confirmed in more than 35 countries.

**Methodology:** We performed a comparative study in the state’s reference hospital for infectious/parasitic diseases, which treats adults with HIV/acquired immune deficiency syndrome (AIDS), between January 2007 and July 2017. The data obtained using this protocol were analyzed using SPSS.

**Principal findings:** In total, 163 patients were evaluated in this study, including 126 patients with coincident VL/HIV and 37 patients with VL alone. Both groups consisted primarily of male patients. The most commonly affected age group was 30–39 years (*p* < 0.001). Fever (*p* < 0.001) and hair loss (*p* = 0.007), which were more common in patients with VL alone, were more common. On hemogram, segmented neutrophils (*p* < 0.0001) were found to be more in the VL/HIV group than in the VL alone group. Additionally, AST and ALT levels differed between the groups (*p* < 0.001). On average, HIV was diagnosed 2.6 years before VL (*p* < 0.001). VL relapse was observed only in the co-infection group (36.5% of cases). Fever (*β* = +0.17; *p* = 0.032) in the first VL/HIV episode was identified as a risk factor for relapse (*R*^2^ = 0.18). The death rate of co-infected patients was 11.1%.

**Conclusion/Significance:** VL/HIV was prevalent among young adults, whereas the median patient age was higher in the VL group. The classic symptomatology of VL was more common in patients not co-infected with HIV, but attention is needed regarding the presence of fever in the first episode of VL as a risk factor for relapse in co-infected patients. No cases of VL relapse occurred in patients without HIV.

**AUTHOR SUMMARY:** Visceral Leishmaniasis (VL) and HIV have maintained increasing rates of populational occurrence in the Northeast region of Brazil, with advancing in rural areas and VL advancing in urban areas. This dynamic explains the increase of co-infection from 0.7% in 2001 to 8.5% in 2012. The state of Maranhão presents a large number of cases in the Northeast Brazil, a region with the majority of patients with this co-infection. The severity presented by these patients, in the observation of frequent relapses and high lethality, is presented in this study, which contributes to epidemiological, clinical, laboratorial, and evolutionary knowledge, trying to demonstrate these in the statistical data. Therefore, it should be pointed out that young adult patients with HIV/AIDS who present with any cytopenia, with or without hepatosplenomegaly and accompanied by fever or not, must be investigated for VL.

## INTRODUCTION

Visceral leishmaniasis (VL) is an infectious parasitic disease transmitted by *Lutzomyia longipalpis*, with *Leishmania infantum chagasi* being the main causative agent in South America [1]. The clinical features of VL may evolve with severity, thus requiring early diagnosis to avoid further complications [2]. In Brazil, 634,051 cases of human immunodeficiency virus/acquired immune deficiency syndrome (HIV/AIDS) had been registered through June 2016 [3]. HIV continues to spread across Brazil, and its incidence in countryside locations has been increasing. This explains its association with other endemic diseases, such as VL, which has also increased in frequency in urban locations, thereby accelerating the clinical evolution of both HIV and VL due to cumulative immunosuppression [4,5]. On the basis of altered epidemiological profiles of both HIV and VL, the risk of exposure to the diseases has grown, with the risk of an HIV-infected person developing VL surging by 100–2320-fold in endemic areas. Moreover, co-infected patients also have reduced responses to therapy, thus resulting in increased risks of recurrence and mortality [6]. The first case of VL/HIV co-infection was reported in 1985, increasing attention in Mediterranean and northern European countries regarding the possibility of this association and the gravity of both diseases [7]. By 2007, 35 countries had already reported cases of VL/HIV co-infection [8]. In Brazil, co-infection was first reported in 1990 in a patient from the Northeast region [9]. In 2015, 257 cases of co-infection VL/HIV were reported in the Western hemisphere, corresponding to a rate of 7.4% of co-infection between all VL cases. Among these, most cases (244) were recorded in Brazil [10]. In 2017, a descriptive and exploratory study that analyzed data from all 21 Brazilian states that had reported cases of VL over a 10-year period identified 1301 cases of VL/HIV co-infection. The researchers noticed an increase in the number of VL/HIV cases over time [11]. Of these states, 4608 cases of VL were recorded in Maranhão, including 256 cases of HIV co-infection [12]. One study performed in the Reference Hospital for Infectious and Parasitic Diseases in Maranhão identified 61 patients with VL/HIV co-infection over a period of 7 years [13]. Considering the high incidence rates of VL in Maranhão and the need to improve knowledge about this pathology, especially epidemiological, clinical, and laboratory data related to VL/HIV co-infection, this study sought to identify particularities in the clinical and laboratory presentation of VL irrespective of its association with HIV. This information may contribute to improving the diagnosis and therapeutic management of patients with VL/HIV co-infection.

## METHODS

### Type of study, location, and population

This comparative study of adults, who are at least 18 years old, was conducted between January 2007 and July 2017 and divided into two steps: (i) a retrospective study of patients diagnosed with VL/HIV co-infection between 2007 and 2015 and (ii) a prospective study of co-infected patients diagnosed between 2016 and 2017. Data were also collected retrospectively from the records of patients with VL without HIV co-infection who were treated at the hospital during the study period. A protocol sheet adapted from the monitoring and control handbook for VL from the Health Ministry, which was named the investigation sheet of death from VL, was used [14]. The sheet was used to record the following variables: sociodemographic (sex, age, occupation, place of residence), epidemiological (place of contamination, other places of residence in the last 12 months), background (other diseases and treatment used), and physical evaluation variables (hydration level, abdominal protrusion, hepatosplenomegaly, swelling); treatment for HIV and leishmaniasis; laboratory examinations (complete blood count, AST, ALT, urea, creatinine, glycemia, CD4 T-lymphocyte count, viral load, bone marrow test); and disease evolution (recurrence, death, hospital release). A case was considered new when the patient had no history or previous record of VL, and deemed recurrent otherwise. The study was conducted at Maranhão’s reference hospital for infectious parasitic diseases, which serves the adult population, including approximately 4640 patients with HIV/AIDS each month.

### Inclusion criteria

The retrospective study included patients with records of VL irrespective of HIV co-infection, containing complete epidemiological, clinical, and laboratory data. Meanwhile, the prospective study included patients with positive serology for HIV and a parasitological diagnosis of VL who were receiving treatment for co-infection at the reference hospital. Meanwhile, the VL group consisted of patients with a parasitological diagnosis of VL and negative serology for HIV who were treated at the reference hospital. All patients included in the prospective study provided written informed consent for participation.

### Diagnosis

#### Visceral Leishmaniasis

*Parasitological*: VL was indicated by the presence of the causative parasite in bone marrow (amastigote forms seen in smears aspirated from bone marrow, stained by Giemsa) [15]. *Serological*: The indirect fluorescent antibody test has been widely used for diagnosing VL, and it is currently provided by Brazil’s Unified Health System. Its sensitivity ranges from 82 to 95%, and its specificity ranges from 78 to 92% [16,17]. Another method used enzyme-linked immunosorbent assay (ELISA), which has sensitivity of 90–100% [18].

#### HIV

HIV was diagnosed using ELISA and the HIV rapid test, which are routinely performed at the reference state hospital following recommendations from the Health Ministry. Other laboratory data used as reference were obtained using the automated system CELL DYN 4000 provided by CEDRO, São Luís, MA, 2002.

#### Statistical analyses

Data were processed using SPSS software version 18.0 (IBM, Chicago, IL, USA). Descriptive statistics was performed by calculating the frequency, average, and standard deviation. The distribution of categorical variables was compared between the groups using the Chi-squared or Fisher’s exact test. The normality of the quantitative variables was measured using the Lilliefors test. Then, an independent Student’s *t*-test (for normally distributed data) and Mann–Whitney’s *U*-test (for non-normally distributed data) were used in the comparative analysis of serum markers between the VL/HIV and VL groups. The coefficient of multiple determination and the coefficient and standard error of each factor included in the model were estimated. For all analyses, a significance level of 5% was adopted.

#### Ethical considerations

The research project was registered at Plataforma Brasil, with ethical committee approval granted by Maranhão’s University Hospital (HUUFMA; date of approval: September 28, 2013, report number: 409.351). Retrospective data were obtained with permission from the ethics committee of HUUFMA and Health Department of the state of Maranhão without obtaining patient consent while maintaining confidentiality regarding the identities of the patients.

## RESULTS

In total, 163 patients were evaluated in this study, including 126 patients with VL/HIV co-infection and 37 patients with VL alone. The demographic and social characteristics of the study groups are described in Table 1. Both groups consisted primarily of male patients. The mean age in the VL/HIV group was 36.9 ± 15 years (range, 19–50), compared to 30 ± 13.4 years (range, 19–66) in the VL group (*p* < 0.001). In both groups, patients were primarily native individuals from Maranhão’s countryside.

**Table 1.** Sociodemographic variables of the studied groups.

| Variable | VL/HIV group<br>(*n* = 126) |  | VL group<br>(*n* = 37) |  | *p* |
| --- | --- | --- | --- | --- | --- |
|  | *n* | % | *n* | % |  |
| Sex |  |  |  |  | 0.768 |
| Male | 113 | 89.7 | 34 | 91.9 |  |
| Female | 13 | 10.3 | 3 | 8.1 |  |
| Age group |  |  |  |  | <0.001* |
| 19–29 years | 19 | 23.0 | 24 | 64.9 |  |
| 30–39 years | 49 | 38.9 | 7 | 18.9 |  |
| 40–49 years | 38 | 30.2 | 3 | 8.1 |  |
| 50+ years | 10 | 7.9 | 3 | 8.1 |  |
| Place of residence |  |  |  |  | 0.213 |
| São Luís | 54 | 42.9 | 11 | 29.7 |  |
| Other municipalities | 72 | 57.1 | 26 | 70.3 |  |
| Occupation |  |  |  |  | 0.560 |
| Self-employed | 26 | 22.4 | 9 | 24.3 |  |
| Agriculture | 25 | 21.5 | 9 | 24.3 |  |
| Construction industry | 21 | 18.1 | 3 | 8.1 |  |
| General services | 14 | 12.1 | 3 | 8.1 |  |
| Housewife | 8 | 6.9 | 2 | 5.4 |  |
| Other | 22 | 19.0 | 11 | 29.7 |  |

VL, visceral leishmaniasis.
\*Statistically significant difference between the groups as determined using the Chi-squared or Fisher's exact test ( $p < 0.05$ ).

The most frequent clinical complaints in the VL group, as showing in table 2, were fever (*p* < 0.001) and hair loss (*p* = 0.007). The physical examination data revealed that weight loss (*p* = 0.003), fever (*p* < 0001), jaundice (*p* < 0.001), hepatomegaly (*p* = 0.029), and splenomegaly (*p* = 0.007) were more frequent in the VL group. In contrast, dyspnea was detected only in the VL/HIV group (*p* = 0.013). The frequency of skin/mucosal pallor was high in both groups (*p* = 0.920).

**Table 2.** Clinical and physical examination data in the two groups.

| Variable | VL/HIV group<br>(*n* = 126) |  | VL group<br>(*n* = 37) |  | *p* |
| --- | --- | --- | --- | --- | --- |
|  | *n* | % | *n* | % |  |
| Clinical complaint |  |  |  |  |  |
| Fever | 66 | 52.4 | 35 | 94.6 | <0.001* |
| Hair loss | 20 | 15.9 | 14 | 37.8 | 0.007* |
| Anorexia | 97 | 77.0 | 32 | 86.5 | 0.307 |
| Diarrhea | 66 | 52.4 | 21 | 56.8 | 0.778 |
| Vomiting | 37 | 29.4 | 16 | 43.2 | 0.166 |
| Bleeding | 27 | 21.4 | 5 | 13.5 | 0.406 |
| Cough | 38 | 30.2 | 10 | 27.0 | 0.871 |
| Dyspnea | 46 | 36.8 | 12 | 32.4 | 0.770 |
| Increased abdominal volume | 38 | 30.2 | 17 | 45.9 | 0.112 |
| Physical symptom |  |  |  |  |  |
| Alopecia | 11 | 8.7 | 4 | 10.8 | 0.747 |
| Weight loss | 66 | 52.4 | 30 | 81.1 | 0.003* |
| Fever | 22 | 17.5 | 25 | 67.6 | <0.001* |
| Jaundice | 22 | 17.5 | 24 | 64.9 | <0.001* |
| Spotting | 11 | 8.7 | 1 | 2.7 | 0.300 |
| Bleeding | 12 | 9.5 | 5 | 13.5 | 0.694 |
| Swelling | 15 | 11.9 | 1 | 2.7 | 0.123 |
| Dyspnea | 17 | 13.5 | 0 | 0 | 0.013* |
| Hepatomegaly | 89 | 70.6 | 33 | 89.2 | 0.029* |
| Splenomegaly | 73 | 57.9 | 31 | 83.8 | 0.007* |
| Adenomegaly | 15 | 11.9 | 4 | 10.8 | 1.000 |
| Abdominal protrusion | 30 | 23.8 | 15 | 40.5 | 0.073 |
| Skin/mucosa color |  |  |  |  | <0.920 |
| Normal color | 27 | 21.4 | 7 | 18.9 |  |
| Pale | 99 | 78.6 | 30 | 81.1 |  |

VL, visceral leishmaniasis.
\*Statistically significant difference between the groups as determined using the Chi-squared or Fisher's exact test ( $p < 0.05$ ).

Table 3 shows the comparative analysis of the serum profiles between groups at the time of admission for VL treatment. The VL/HIV group exhibited a higher frequency of lymphocytopenia when compared to the VL group (*p* < 0.001). Abnormal AST and ALT levels were observed more commonly in the VL group (*p* < 0.001). Most co-infected patients had CD4 T-lymphocyte counts of less than 200 cells/mm^3^ and viral loads of 50–9999 copies/mL, although the diagnosis of HIV came, in average, 2,6 years sooner than the one of VL in 81 patients (64.3%, *p* < 0.001).

**Table 3.** Laboratory data of the two groups.

| Variable | VL/HIV group<br>(*n* = 126) |  | VL group<br>(*n* = 37) |  | *p* |
| --- | --- | --- | --- | --- | --- |
|  | *n* | % | *n* | % |  |
| Red blood cells (million/mm<sup>3</sup>) |  |  |  |  | 1.000 |
| <4.6 | 118 | 93.6 | 35 | 94.6 |  |
| 4.6–6.2 | 8 | 6.4 | 2 | 5.4 |  |
| Hemoglobin (g/dl) |  |  |  |  | 0.425 |
| <14.0 | 120 | 95.2 | 34 | 91.9 |  |
| 14–17 | 6 | 4.8 | 2 | 8.1 |  |
| Hematocrit (%) |  |  |  |  | 0.352 |
| <40 | 119 | 94.4 | 37 | 100 |  |
| 40–54 | 7 | 5.6 | 0 | 0 |  |
| Leukocytes (1000/mm<sup>3</sup>) |  |  |  |  | 0.224 |
| <5000 | 101 | 80.2 | 34 | 91.9 |  |
| 5000–10,000 | 22 | 17.5 | 3 | 8.1 |  |
| >10,000 | 3 | 2.3 | 0 | 0 |  |
| Lymphocytes (%) |  |  |  |  | <0.001* |
| <20 | 30 | 23.8 | 3 | 8.1 |  |
| 20–40 | 79 | 62.7 | 18 | 48.7 |  |
| >40 | 17 | 13.5 | 16 | 43.2 |  |
| Platelets (1000/mm<sup>3</sup>) |  |  |  |  | 0.083 |
| <150,000 | 79 | 62.7 | 28 | 75.7 |  |
| 150,000–400,000 | 45 | 35.7 | 7 | 18.9 |  |
| >400,000 | 2 | 1.6 | 2 | 5.4 |  |
| Urea (mg/dl) |  |  |  |  | 0.449 |
| 10–50 | 105 | 83.3 | 33 | 89.2 |  |
| >50 | 21 | 16.7 | 4 | 10.8 |  |
| Creatinine (mg/dl) |  |  |  |  | 0.820 |
| <0.4–1.2 | 110 | 87.3 | 31 | 83.8 |  |
| >1.2 | 16 | 12.7 | 6 | 16.2 |  |
| AST |  |  |  |  | <0.001* |
| 12–46 | 76 | 60.3 | 10 | 27.0 |  |
| <46 | 50 | 39.7 | 27 | 73.0 |  |
| ALT |  |  |  |  | <0.001* |
| 3–500 | 102 | 80.9 | 16 | 43.2 |  |
| <50 | 24 | 19.1 | 21 | 56.8 |  |
| CD4 T-lymphocytes<br>(cells/mm<sup>3</sup>) |  |  |  |  |  |
| <200 | 98 | 77.8 | – | – |  |
| ≥200 | 28 | 22.2 | – | – |  |
| Viral load (copies/mL) |  |  |  |  |  |
| <50 | 19 | 15.1 | – | – |  |
| 50–9999 | 68 | 54.0 | – | – |  |
| 10,000–100,000 | 13 | 10.3 | – | – |  |
| >100,000 | 26 | 20.6 | – | – |  |

VL, visceral leishmaniasis.
\*Statistically significant difference between the groups through as determined using the Chi-squared or Fisher's exact test ( $p < 0.05$ ).

The distribution of variables related to treatment is shown in Table 4. The most commonly used medicine in the VL/HIV group was liposomal amphotericin B (Amb-L), whereas it was *N*-methyl meglumine antimoniate (Sb^v^) in the VL group (*p* < 0.001). The treatment time for VL was no more than 10 days in most patients in the co-infection group (*p* < 0.001). In addition, 112 patients (88.9%) in the VL/HIV group remained hospitalized for more than a month, whereas no patients in the VL group were hospitalized (*p* < 0.001). Furthermore, relapse occurred only in the co-infection group (*p* < 0.001). Concerning antiretroviral treatment, highly active antiretroviral therapy was administered to all analyzed patients; most commonly a nucleoside reverse transcriptase inhibitor in combination with a non-nucleotide reverse transcriptase inhibitor or protease, including tenofovir + lamivudine in 113 (89.7%) patients, efavirenz in 54 patients (42.9%), and lopinavir/ritonavir in 52 patients (41.7%). Moreover, relapse (46 cases, 36.5%) and death (14, 11.1%) occurred only in the VL/HIV group.

**Table 4.** VL treatment in the two groups.

| Variables | VL/HIV group<br>(*n* = 126) |  | VL group<br>(*n* = 37) |  | *p* |
| --- | --- | --- | --- | --- | --- |
|  | *n* | % | *n* | % |  |
| VL treatment |  |  |  |  | <0.001* |
| Liposomal amphotericin B | 86 | 68.2 | 2 | 5.4 |  |
| Amphotericin B | 24 | 19 | 1 | 2.7 |  |
| *N*-methyl meglumine antimoniate | 8 | 6.4 | 34 | 91.9 |  |
| More than one drug | 8 | 6.4 | 0 | 0 |  |
| Duration of treatment |  |  |  |  | <0.001* |
| Up to 10 days | 91 | 72.8 | 0 | 0 |  |
| 11–30 days | 27 | 21.6 | 31 | 83.8 |  |
| More than 30 days | 7 | 5.6 | 6 | 16.2 |  |
| Duration of hospitalization |  |  |  |  | <0.001* |
| None | 0 | 0 | 37 | 100 |  |
| Less than 1 month | 14 | 11.1 | 0 | 0 |  |
| 1–7 months | 109 | 86.5 | 0 | 0 |  |
| More than 6 months | 3 | 2.4 | 0 | 0 |  |
| VL relapse |  |  |  |  | <0.001* |
| Yes | 46 | 36.5 | 0 | 0 |  |
| No | 80 | 63.5 | 39 | 100 |  |
| Death |  |  |  |  | <0.001* |
| Yes | 14 | 11.1 | 0 | 0 |  |
| No | 0 | 0 | 0 | 0 |  |

VL, visceral leishmaniasis.
\* Statistically significant difference between the groups as determined using the Chi-squared or Fisher's exact test ( $p < 0.05$ ).

## DISCUSSION

Visceral Leishmaniasis, an endemic disease in the northern and northeastern regions of Brazil, has changed in its epidemiological characteristics with the spread of HIV infection toward rural areas and coincidence of the diseases. In patients infected with HIV, VL can induce greater immunosuppression and stimulate viral replication, leading to a more aggravated condition and greater risks of relapse and death [19, 2, 20, 21].

VL/HIV co-infection displayed a heterogeneous casuistry among the Brazilian regions, but its incidence appears to be increasing across the country. The rates of co-infection according to Lindoso *et al.* were 0.7% in 2001 and 8.5% in 2012 [22].

Between 2007 and June 2017, data were collected for 127 patients with VL/HIV co-infection at the reference hospital where the study was performed. This cohort represented nearly 50% of all reported cases in the state of Maranhão through December 2016 [12]. The finding of a higher incidence of VL in males in both VL with HIV and VL alone is supported by other findings, including epidemiological data from the Health Ministry, which reported that male patients accounted for 91.6% of all cases [2,23,24]. Similarly, the finding that the mean age was lower in the VL group matches data reported by Hurissa *et al.*, who found that patients with VL alone had a mean age of 23.5 years. However, Cota *et al.* found no statistical difference in age between the VL/HIV and VL groups in a study of 90 patients [24,25].

The majority of diagnosed patients resided in Maranhão’s countryside, as corroborated by Furtado, who analyzed the residences of patients with VL in the state of Maranhão based on the National System for Reporting Harms, applying a method for risk identification, between 2000 and 2009. This analysis found that the incidence rates were higher in the cities of Caxias, Imperatriz, Presidente Dutra, Codó, and Barra do Corda, with attention also being devoted to the appearance of cases in countryside cities in which no VL case had been previously reported [26]. By observing the internalization of VL, it is possible to perceive the juxtaposition that occurs between these VL infections found in people living with HIV an adequate environment to develop [27].

In line with our findings, Cota *et al.* described fever and hepatosplenomegaly as the most frequent complications in patients with VL alone. In contrast, Hurissa *et al.* found typical VL complications in both studied groups [24,25]. In their study, fever was observed more frequently in patients with VL than those with VL/HIV co-infection, group that presented this symptom in 52,4% of the patients, whereas this symptom was present in 100% of patients diagnosed in the state of Mato Grosso do Sul between 2000 and 2006 and 92% of patients diagnosed in the state of Ceará in 2015 [23,28]. Different from many Brazilian studies, which reported the presence of splenomegaly followed by fever, weight loss, and asthenia, in 81 patients with VL/HIV co-infection in Ceará [29] and another study that identified weight loss, weakness, fever, and hepatosplenomegaly as the most common physical changes in 65 patients with VL/HIV co-infection [13], the most common symptoms in this study were skin pallor, hepatomegaly, and splenomegaly, followed by fever and weight loss. In a systematic review of the epidemiological, clinical, and laboratory aspects of VL associated with HIV, the authors noted that the 10 most reported clinical manifestations in 15 studies were skin pallor, splenomegaly, fever, weight loss, hepatomegaly, couch, diarrhea, bleeding, asthenia, and jaundice [30].

Pancytopenia was observed in both studied groups, in contrast to the findings of Cota *et al.* and Hurissa *et al.*, who observed more pronounced thrombocytopenia in immunocompetent patients [24,25]. Pancytopenia related to HIV can be multifactorial due to direct effects of the virus, ineffectiveness of hematopoiesis, existence of an infiltrating disease in bone marrow, nutritional deficiencies, peripheral destruction, and toxic medication effects [31]. In contrast, hematological alterations caused by VL are well known, as amastigote forms proliferate in the mononuclear phagocytic system, mainly in the spleen, liver, and bone marrow, resulting in disorders in phagocytic organs and producing hematological alterations. Tavora *et al.* observed pancytopenia in the majority of patients with VL/HIV co-infection [28,32]. The finding of increased creatinine levels in less than 20% of cases in both groups was also reported by Mahajan *et al.*, who observed an increase in creatinine levels in 16 patients (5% of the patients with VL-HIV co-infection). Sinha *et al.*, who analyzed 49 patients with VL/HIV co-infection, found a mean creatinine level of 0.9 mg/mL. The increase in transaminase levels observed in some patients with VL/HIV co-infection [33,34,35] was also found in the present study. However, we should note that such an increase was higher in patients with VL alone, as additionally reported by De Souza *et al.* and Cota *et al.* [25,36].

CD4 T-lymphocytes were quantified by several authors who emphasized, similar to our study, the presence of immunosuppression with counts lower than 200 cells/mm^3^ in a large portion of co-infected patients at the time of VL diagnosis [23,34,35,37,38]. However, we emphasize that the diagnosis of HIV having been performed before VL in most co-infected patients, these patients should be immunologically less suppressed since they would be taking antiretroviral, but most of them had a viral load above 50 copies. This may have been due to non-adherence to antiretroviral therapy, virological failure or worsening caused by VL infection, as it been shown that co-infected patients may have increased immunosuppression and stimulation of viral replication [8].

Treatment for VL was administered according to the Health Ministry protocol, with AmB-L being the drug of choice for patients with VL/HIV co-infection starting in 2015, whereas Sb^v^ was used to treat patients with nonserious VL [2]. Brazilian authors have reported treatments that reflect guidance from the Health Ministry, drawing attention to the adverse effects of AmB such as kidney toxicity and reactions during infusion. Toxicities linked to Sb^v^ administration were also reported, in some cases requiring substitution with AmB. Some of the reports cited pancreatitis and cardiotoxicity [23,25,36,39]. The treatment duration for VL varied according to the therapeutic response using the criteria adopted by the Health Ministry, which call for a minimum duration of 10 days [19]. However, we observed therapeutic failure, even after increasing the duration of AmB-L administration, making supplemental therapy with Sb^v^ necessary (data not cited). AmB-L has been considered promising for the treatment of VL/HIV co-infection since 1999, when it was approved by the FDA [40]. However, although it is associated with fewer side effects than AmB, VL/HIV co-infection remains to be associated with high rates of recurrence, as noted in the current study. Lindoso *et al.* noted that no standard therapeutic regimes have been developed for VL in the United States [22]. Cota *et al.* reported recurrence and mortality rates of 37 and 8.7%, respectively, in a group of 46 patients with VL/HIV co-infection, compared to those of 2.5 and 4.7%, respectively, in 44 patients with VL. In a sample of 128 patients with VL, Oliveira reported recurrence and death rates of 7.8 and 4.7%, respectively. This result differs from that of the present study, in which no case of recurrence or death was reported among patients with VL mono-infection versus a mortality rate of 11.1% among patients with VL/HIV co-infection [25,41].

This study had several limitations. Because of the use of secondary data, mainly in the retrospective study, incomplete data were obtained in some examinations. Another limitation was difficulty in analyzing some laboratory data, which precluded comparative analyses between the two groups, although we observed that the co-infection group patients had a lower rate of positivity in serological or rapid tests than the group without co-infection. Another limitation was the lower number of patients with VL alone. This finding was due to the lack of information in the medical records of these patients, as most of them were diagnosed in this referral hospital and subsequently referred for treatment at locations closer to their places of residence. On the contrary, the patients with VL/HIV co-infection in this study represented almost half of the 256 cases of co-infection in the state of Maranhão over the period of 2007–2016. We included patients only for whom complete data were available, increasing the statistical power of the analyses.

In summary, our study found that VL is a public health issue in northeastern Brazil, especially in Maranhão, where its incidence is increasing. The severity of the disease is greater due to co-infection with HIV, which modifies the epidemiology and clinical presentation of VL. Thus, the classical triad of fever, paleness, and hepatosplenomegaly was not expected. We also concluded that treatment remains challenging, despite advances in antiretroviral treatment, so a definitive response to treatment for VL cannot be predicted when the disease is associated with HIV.

## SUPPORTING INFORMATION

S1 Checklist: STROBE Checklist

## Acknowledgments

We thank Fundação de Amparo a Pesquisa do Maranhão (FAPEMA), Presidente Vargas Hospital, and Maranhão Health Secretariat.

